# Short-term stimulation of collective cell migration in tissues reprograms long-term supracellular dynamics

**DOI:** 10.1101/2021.07.27.453602

**Authors:** Abraham E. Wolf, Matthew A. Heinrich, Isaac B. Breinyn, Tom J. Zajdel, Daniel J. Cohen

## Abstract

The ability to program collective cell migration can allow us to control critical multicellular processes in development, regenerative medicine, and invasive disease. However, while various technologies exist to make individual cells migrate, translating these tools to control myriad, collectively interacting cells within a single tissue poses many challenges. For instance, do cells within the same tissue interpret a global migration ‘command’ differently based on where they are in the tissue? Similarly, since no stimulus is permanent, what are the long-term effects of transient commands on collective cell dynamics? We investigate these questions by bioelectrically programming large epithelial tissues to globally migrate ‘rightward’ via electrotaxis. Tissues clearly developed distinct rear, middle, side, and front responses to a single global migration stimulus. Furthermore, at no point post-stimulation did tissues return to their pre-stimulation behavior, instead equilibrating to a third, new migratory state. These unique dynamics suggested that programmed migration resets tissue mechanical state, which was confirmed by transient chemical disruption of cell-cell junctions, analysis of strain wave propagation patterns, and quantification of cellular crowd dynamics. Overall, this work demonstrates how externally driving the collective migration of a tissue can reprogram baseline cell-cell interactions and collective dynamics, even well beyond the end of the global migratory cue, and emphasizes the importance of considering the supracellular context of tissues and other collectives when attempting to program crowd behaviors.

---

Collective cell migration is essential for many multicellular organisms and underpins diverse processes, spanning organ development to cancer invasion (1–3). Therefore, externally directing collective cell migration should allow us to not only better understand these processes, but also to formally program and engineer them for applications in regenerative medicine and tissue engineering (4–8). Programming cell migration is increasingly feasible with myriad emerging approaches spanning chemotactic devices (cells migrate along chemical gradients) (8–13), light-induced, optogenetic modulation of signaling pathways (14–16), and bioelectric programming of cell migration through electrotaxis (cells migrate along electrical gradients) (17–22). However, a key challenge is understanding how global commands, such as ‘All Cells Migrate Rightward’, play out in a tissue context with 10,000+ interacting cellular agents at macro-scale.

It is well established that cell-cell interactions can cause a different stimulus response to that observed with single cells. For instance, studies have demonstrated both that cellular groups often better follow chemical (11) and electrical migration cues (23) than do single cells, while overly strong cell-cell interactions and cell-cell coupling can actually compete with external migratory commands leading to adverse outcomes (7, 24). Motivated by these considerations, we investigated two key factors for externally directing collective cell migration: [1] how does where a cell is located within a tissue modulate its response to a global command; [2] does following this global command change the longer term behavior of the tissue after the command is removed?

From huddles of penguins on the ice, to schools of fish, and migrating clusters of cells, it is increasingly clear that the location of an agent (group member) within a group strongly affects how that agent behaves (17, 25–29). In the tissue context, a growing body of work has revealed how gradients of mechanical tension underlie the growth and motion of highly collective tissues, such as the epithelia lining our organs (30–32), and naturally give rise to behavioral differences between boundaries, edge zones, and bulk of a tissue (33, 34). Such behavioral zones can emerge even in groups of loosely coupled cells migrating along chemical gradients (35). This compartmentalization of behaviors within a group is referred to as ‘supracellularity’ (35), and emphasizes how collectives are often best characterized as a whole unit. Applying this supracellular framework here provides an analytical basis to understand how, when, and where cells within a single tissue respond to the same global migratory command.

Additionally, since no migratory command or stimulus is permanent, it is critical to understand the collective response of a cellular group or tissue not only during a global migratory stimulation, but also after removal of the stimulus. Considering the whole process, from initial entrainment of cells to eventual relaxation, can reveal the longer term consequences of externally driving collective cell behaviors, and can detect a collective ‘memory’ of the stimulation event. To date, such cellular memory of migratory stimuli has been studied primarily with single cell relaxation after exposure to chemotactic or electrotactic cues (9, 19). After a chemotactic gradient is removed, internal signaling such as Ras activity may relax over a 30 second scale (36), while front-rear cell polarity and overall directionality decay within ~10 minutes post-stimulation (9, 10). In electrotaxing single cells, directionality has been reported to persist over a ~ 15-60 minute period post-stimulation (19, 37). However, the collective nature of a tissue relative to single cell studies means tissues will necessarily exhibit different relaxation and reprogramming dynamics in response to a global migration command, and investigating the spatiotemporal response of the tissue to external commands will go a long way to improving both our understanding of collective cell behaviors, and our ability to more effectively program and augment large-scale tissue behaviors for tissue engineering.

To investigate these questions, we needed both a model system and an appropriate migration stimulus. We began with a gold-standard collective cell migration model—the MDCK renal epithelium, which is one of the most well-characterized collective cell systems to date (38–40). MDCK epithelia are readily grown to centimeter scale, and exhibit robust outward migration and canonical ‘scratch wound’ healing dynamics (33, 34, 41), enabled by strong cell-cell coupling mediated by E-cadherin and other junctional complexes (17, 31, 42). We next needed a compatible stimulus that could precisely and globally induce directional migration across the whole epithelium. Chemotaxis and electrotaxis are two of the best characterized directional migration candidates. However, while chemotaxis is potent and well-understood in certain single cells and small cluster models, many cell types and model systems lack chemotactic pathways or receptors (including MDCK cells), so chemotaxis needs to be engineered and fine-tuned in cells that do not natively chemotax (8). Electrotaxis, by contrast, arises naturally in most cell types and is conserved across many species (19, 43–46). Electrotaxis shares certain key signaling pathways with chemotaxis (notably PI3K and TORC2) (46), and induces directional migration based on naturally occurring ionic current gradients (*O(1 V/cm)*) (47, 48) formed during tissue perturbations (e.g. morphogenesis, healing). These fields apply weak electrophoretic and electro-osmotic forces to membrane receptors, inducing receptor aggregation and activation, leading to cell polarity and migration (19, 49). That electrotaxis affords programmatic control of collective cell migration with uniform and near-instantaneous stimulus of the whole tissue (18, 24, 50, 51), without requiring additional reagents or cellular modifications, made it an optimal ‘command’ stimulus for this study.

We then used automated phase contrast microscopy to capture supracellular patterns and relaxation dynamics of epithelial tissues exposed to a transient electric field inducing ‘rightward’ electrotaxis. We first quantify how the various edges (leading, trailing, top, bottom) of a ‘rightward’ migrating tissue exhibit distinct behaviors, with particular emphasis on apparent elastic recoil in these zones once stimulation is removed. These responses were markedly different from how the bulk center of tissues behave, where we observed long-term directional ‘memory’ long after stimulation was stopped. We link this memory to a resetting of tissue mechanical state that must occur in order to enable collective migration of mechanically coupled cells, which we validate by characterizing strain wave dynamics before, during, and after stimulation. Finally, we demonstrate how driven collective migration ultimately reprograms the underlying cell-cell correlations in a lasting manner, meaning that a post-stimulation tissue differs markedly from its unstimulated, control sample counterpart.

## Results

### Programmed collective migration alters supracellular migration patterns in tissues

To generate the core data needed to explore supracellularity and programmed migration, we captured the complete step response of large epithelia to global bioelectric migration commands (Fig. 1A-C, Movie 1 compares control and stimulated tissues). This required continuous time-lapse imaging of arrays of 5×5 mm MDCK epithelial monolayers over three contiguous periods: control (1 h), stimulation ON (3 h), and stimulation OFF (6 h). Movie 2 shows closer views of local cellular responses during this process in control and stimulated tissues. To induce electrotaxis, we built custom electro-bioreactors around these tissue arrays similar to our SCHEEPDOG platform (7, 18), which allowed us to provide continuous media perfusion and a computer-controlled, uniform, ‘rightward’ electrical cue (3 V/cm field) to all tissues in the bioreactor. A schematic of our bioreactor and representative epithelium highlighting our analysis zones are presented in Figs. 1A-B.

**Fig. 1.**
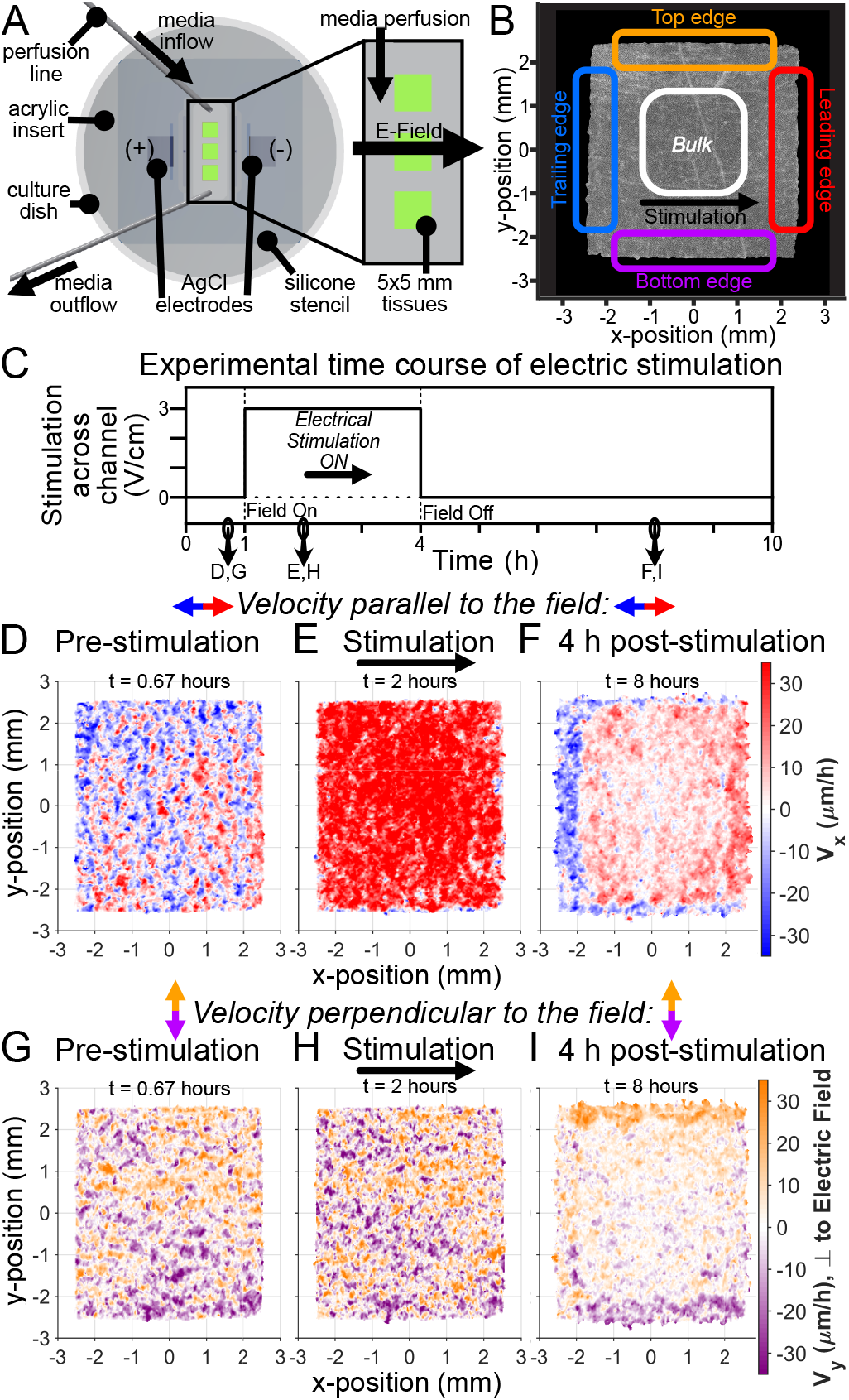
Collective electrotaxis step response induces a spatiotemporal effect in epithelial tissues persisting long after stimulation. *(A)* Diagram of electro-bioreactor providing electrical stimulation in the *x*-axis and perfusion of fresh media in the *y*-axis through the chamber containing three 5 × 5mm tissues. *(B)* Phase contrast image of a single MDCK epithelial tissue; annotations denote specific regions analyzed throughout the paper. *(C)* Trace of electrical stimulation step response stimulus: 1 h unstimulated, 3 h ‘rightward’ stimulation, and 6 h post-stimulation. *(D-F)* Single tissue heatmaps of velocity in the *x*-direction, aligned with the direction of electric field (red). Note that *E* indicates large increase in rightward collective motion during stimulation, while *F* indicates a new relaxation behavior hours after stimulation. See Fig. S1. for corresponding control tissue heatmaps. *(G-I)* Analogous heatmaps of velocity in the *y*-axis, perpendicular to the direction of electric field. Note again the new pattern long after stimulation ended in *I*, despite minimal apparent changes during stimulation in *H*.

To begin, we measured collective migration patterns using particle image velocimetry on phase contrast images (PIV; see Methods) to map the spatial velocity profile across the tissue and throughout the experiment (Fig. 1C). We report velocities parallel (*V_x_*, Figs. 1D-F) and perpendicular (*V_y_*, Figs. 1G-I) to the direction of stimulation (full heatmaps presented in Movie 3 and Fig. S1). In the direction parallel to the field, we saw clear progression of the velocity field, from relatively disordered during the control period (Fig. 1D), to higher magnitude and well-aligned with the field during stimulation (Fig. 1E), and finally to a distinct relaxation phase post-stimulation (Fig. 1F). In the direction perpendicular to the field (Figs. 1G-I), we observed no obvious change during stimulation, but a similar post-stimulation state to Fig. 1F, except parallel to the *y*-axis.

While directed collective migration during stimulation is expected with tissue electrotaxis (4, 17, 18, 50), the distinct migratory patterns long after stimulation had ceased, and the fact that these patterns did not resemble any previous state were surprising and previously unreported. Furthermore, the migration patterns visible in post-stimulation heatmaps, where the bulk zone retained a ‘memory’ of rightward migration (red center, Figs. 1E-F) while the perpendicular and trailing edges appeared to ‘recoil’ and migrate opposite to the prior command (blue edges, Fig. 1F), suggest a long-term reprogramming of the supracellular dynamics. These compartmentalized behavioral zones indicated where to focus our further analyses.

### Tissue boundaries compete with external migration commands and rapidly recover post-stimulation

We separately analyzed the different edge and bulk zones to better understand how a tissue parses a global migration command, beginning with the leading and trailing edge dynamics, as edge outgrowth is very well characterized in unperturbed epithelia (33, 34, 39). Here, we found that while both the leading and trailing edges obeyed the migration command and exhibited greater ‘rightward’ migration speed during stimulation, they very rapidly equilibrated back to control growth speeds in <30 minutes post-stimulation (Fig. 2A; see also Figs. 1D-F). Interestingly, the trailing edge exhibited a transient overshoot of outgrowth speed that was quickly damped within 30 minutes post-stimulation, suggestive of the viscoelastic nature of epithelial tissues (52, 53). To resolve how individual cellular agents behaved in the edge zone, we performed single-cell tracking of all cells within 150 *μ*m of the trailing edge and overlaid cell trajectories in a ‘hairball’ plot (Fig. 2B; see Methods). This clearly demonstrates both directed migration following the stimulus (pink line segment of Fig. 2B), and how cells performed rapid ‘U-turns’ immediately post-stimulation and quickly equilibrated to control edge dynamics.

**Fig. 2.**
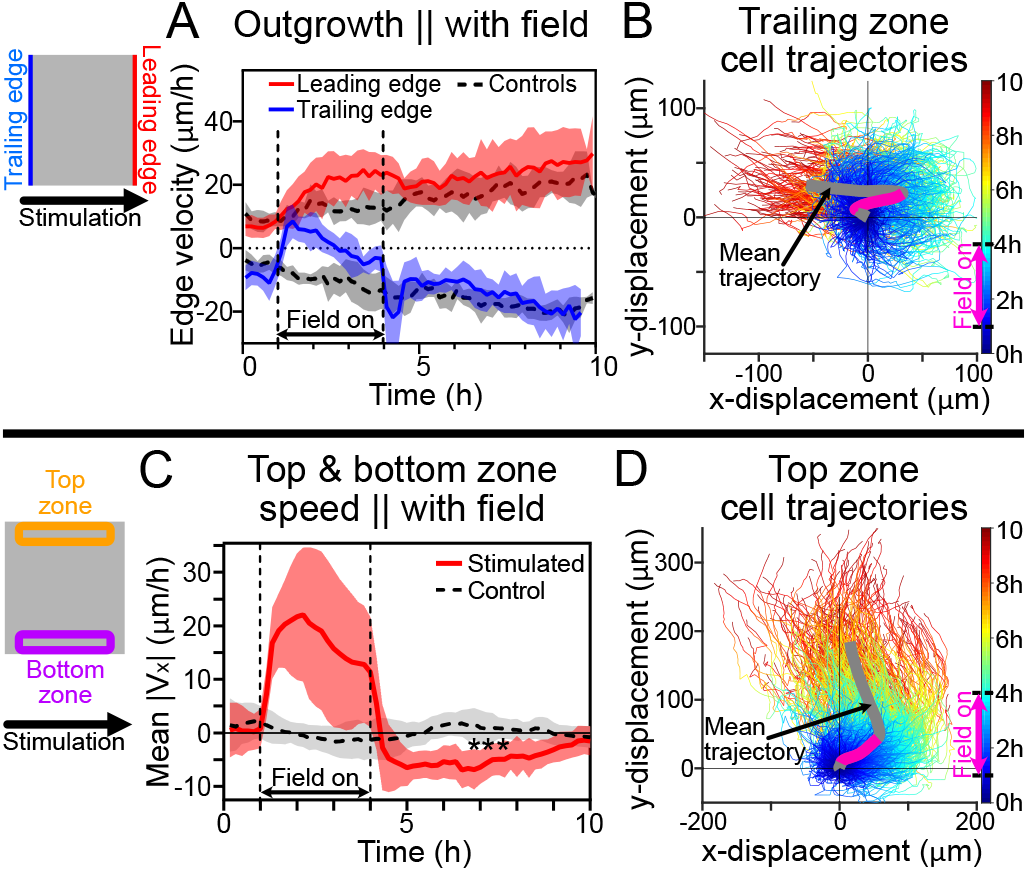
Edges parallel and perpendicular to the field direction displayed distinct behaviors. Cartoons indicate analysis zones. *(A)* Edge outgrowth speed (positive is rightward motion). Control edges (gray) exhibit steady outwards motion, while stimulated edges exhibit perturbed outgrowth during stimulation with rapid post-stimulation recovery. *(B)* Cell trajectories from the trailing zone. Lines color-coded by time (see color bar). Mean trajectory of all tracked trailing zone cells indicated with bold gray line; magenta overlay indicates period of active stimulation. All trajectories were plotted as positive final y-displacements. See Movie 3. *(C)* Average speed in the direction of the electric field in top and bottom edge zones. Note strong increase in rightward motion in these zones during stimulation and rapid over-correction post-stimulation (anti-parallel motion). ***: *p* = 0.0002 between stimulated and control tissues over 6-10 h. *(D)* Cell trajectories from top edge zone. Note again recoiling motion post-stimulation. For panels *A,C*, shading indicates standard deviation across *N_control_* = 6, *N_stimulated_* = 9; panels *B,D* represent single tissues, where zones exclude 1.5 mm on each side for edge effects.

More surprising was the behavior of the top and bottom edges of the tissue. As indicated in Fig. 1, these edge zones largely followed the electrotactic command during stimulation (Figs. 1E,H; Fig. S1), but then exhibited retrograde motion post-stimulation as the epithelium re-established conventional expansion into the free surrounding space, qualitatively reminiscent of elastic recoil. Again analyzing a 150 *μ*m zone at the top and bottom edges, we plotted the ensemble migration velocity (from PIV) in the direction of the stimulus (Fig. 2C; see also Figs. 1,1S). While cells in both top and bottom border zones migrated in the programmed ‘rightward’ direction and increased speed during stimulation, their post-stimulation dynamics exhibited strong, persistent ‘leftward’ migration over several hours (Figs. 2C and S2, *p* = 0.0002 vs. control over 6-10 h; see Methods). A hairball plot of cells in the ‘top’ zone (Fig. 2D) again demonstrates initial entrainment to the electrical command (pink line segment of Fig. 2D) followed by a distinct ‘recoil’ post-stimulation as the tissue re-equilibrates to the conventional outward migration. The dynamics of this process are highlighted in Movie 4 showing a tracked group of cells near the top boundary (see Methods). Overall, these distinct edge dynamics demonstrate that external commands compete with the strong outward migration program of epithelial tissue edges, resulting in a local tug-of-war, eventually won by the outward migration program post-stimulation. Further, this apparent recoil in the non-leading edges is suggestive of elastic recovery after removal of a stress caused by electrical stimulation, consistent with prior work indicating that electrotaxis notably elevates intercellular stresses in the bulk zone of a tissue (54).

### Cells in the tissue center exhibit a memory of the prior migratory command

In contrast to the distinct edge recoil behaviors, the bulk of the tissue relaxed far more gradually and appeared to be the primary source of migratory ‘memory’. To best emphasize this, we analyzed the bulk zone using standard ensemble analyses for directed collective cell migration—the directionality order parameter and mean speed (17, 23, 24, 55), both plotted over the entire experimental time course. Here, we define the directionality order parameter as

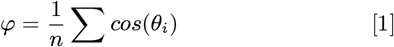

where *θ* is the angle between a PIV velocity vector and the direction of stimulation, the ‘command’ direction unit vector. *φ* ranges from [-1,1], with −1 and 1 indicating cells moving exactly antiparallel and parallel to the command direction, respectively. *φ* = 0 indicates no net directionality at the ensemble level, as is the case for the control data shown (Fig. 3A, black dashed line). In stimulated tissues (Fig. 3A, red line), the directionality dynamics clearly resolve the fast transition from control (*φ* ~ 0) to stimulated (*φ* ~ 1), followed by a much slower relaxation of the tissue migration direction post-stimulation. In contrast, while speed in the direction of stimulation, |*V_x_*|, increased nearly 5× during electrotaxis (Fig. 3B), this effect rapidly decayed to a level that was actually lower than the speed of an equivalent control tissue (Fig. 3B, *p* = 0.003 for 6-10 h; see Methods).

**Fig. 3.**
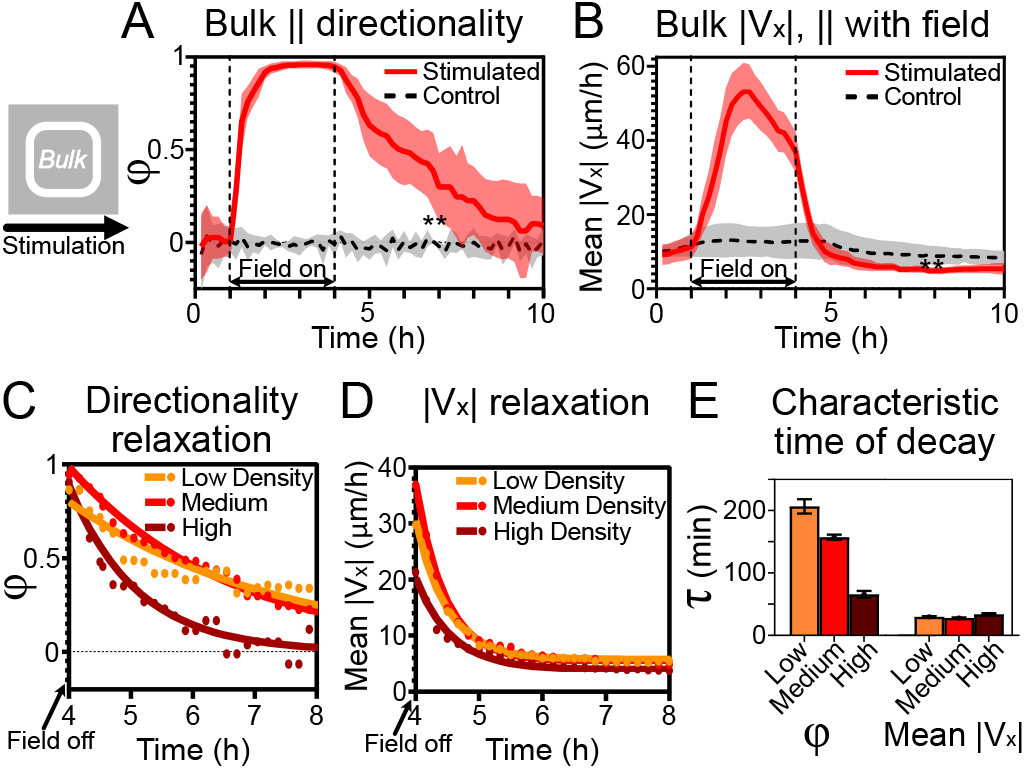
Tissue centers show migratory memory post-stimulation. All data here are captured from the tissue bulk region (see cartoon). *(A)* Directionality order parameter *φ* comparison between unstimulated (dashed gray) and stimulated (red) tissues. Note sustained directionality even post-stimulation. **: *p* = 0.003 between mean values of stimulated and control tissues over 6-10 h. *(B)* Average horizontal speed (parallel to field) during stimulation. Note both strong rise during stimulation and sharp fall post-stimulation to below control levels. **: *p* = 0.003 over 6-10 h. *(C-D)* Exponential decay modeling of post-stimulation relaxation dynamics *τ* for low, medium and high density tissues (see Methods). Raw data in *C-D* indicated by dots; solid curves represent best fits. *(E)* Quantification of characteristic timescales with respect to cell densities for directionality (left) and horizontal speed (right). Note slow directionality relaxation and inverse relation between density and directionality relaxation, which is not true for speed relaxation. For panels *A-B*, *N_control_* = 6, *N_stimulated_* = 9; shading indicates standard deviation; *C-E* include N=5, 9, 4 for low, medium, and high density tissues, respectively. In all panels, tissues’ center 2 × 2 mm were analyzed.

To quantify this difference in relaxation times between ensemble directionality and speed, we defined these timescales *τ* by fitting the data in the post-stimulation phase to an exponential decay model (Figs. 3C-D; see Methods). Here, the characteristic relaxation time *τ* for the directionality order parameter (~ 160 min) was greater than 5-fold longer than for speed (~ 30 min), as shown in Figs. 3C-E (red). To put this in context, these epithelial cells lost ~80% of their previous speed within 20 minutes (losing ~1 *μ*m/min), yet maintained an imperative and ‘memory’ to continue migrating ‘rightward’ for many hours (until *φ* relaxed closer to 0). To add further biophysical context, and motivated by the role cell density plays in modulating collective cell migration and tissue mechanics (33, 56, 57), we repeated these relaxation studies in the tissue bulk at two additional densities, the lower bound being the minimum density that maintained confluence, ~ 2000 cells/mm^2^ (Figs. 3C-E, orange), and the upper bound, ~ 4500 cells/mm^2^ (Figs. 3C-E, brown), sufficient to geometrically limit migration (see Methods) (58). Relaxation dynamics versus density are shown in Figs. 3C-E. While speed relaxed independently from density, migration directionality relaxation times were inversely correlated with density (Fig. 3E), ranging from ~ 3 h (Fig. 3E, low density) to ~ 1 h (Fig. 3E, high density). We hypothesize that this trend may be due to increased cell-cell coupling at higher densities (57–59), or the effects of contact inhibition of locomotion and the onset of jamming at particularly high densities that would disrupt persistent cell motion (3, 60, 61). This slow relaxation of directionality in the bulk is quite distinct from the rapid post-stimulation response in the tissue edges (Fig. 2), perhaps suggesting a decoupling of the bulk from the edges. To investigate this further, we next characterized interplay between edge and bulk dynamics.

### Halting bioelectric stimulation resets the mechanical state of the tissue

To better capture the supracellular dynamics’ coupling behaviors from the edges to the bulk, we generated average kymographs of the directionality order parameter, *φ*, across the entire width of the tissue (averaged vertically; see Methods) over the entire experimental duration (Figs. 4A-C, red again indicates rightward motion). We first established a control case as a reference, Fig. 4A, demonstrating canonical, steady outward tissue expansion in both directions. By contrast, while electrical stimulation induced an overwhelmingly ordered response during the stimulation period (Fig. 4B, red center in 1-4 h; see also Fig. 1E), the post-stimulation dynamics revealed pronounced, inward traveling waves of migration mobilization that nucleated at the outer edges of the tissue and propagated inward at an apparently constant rate (‘triangular regions’ marked with dotted lines in Fig. 4B). Such waves also clearly emerged in post-stimulated tissues from the top and bottom edges in *V_y_* kymographs (Fig. S3). Generally, inward traveling epithelial waves have previously been reported as a natural consequence of releasing a confluent epithelium from confinement and allowing it to grow outward (as in Fig. 4A) (62–64). However, the post-stimulation effect observed here was far stronger than what we observed in our control tissues (compare Figs. 4A,B).

**Fig. 4.**
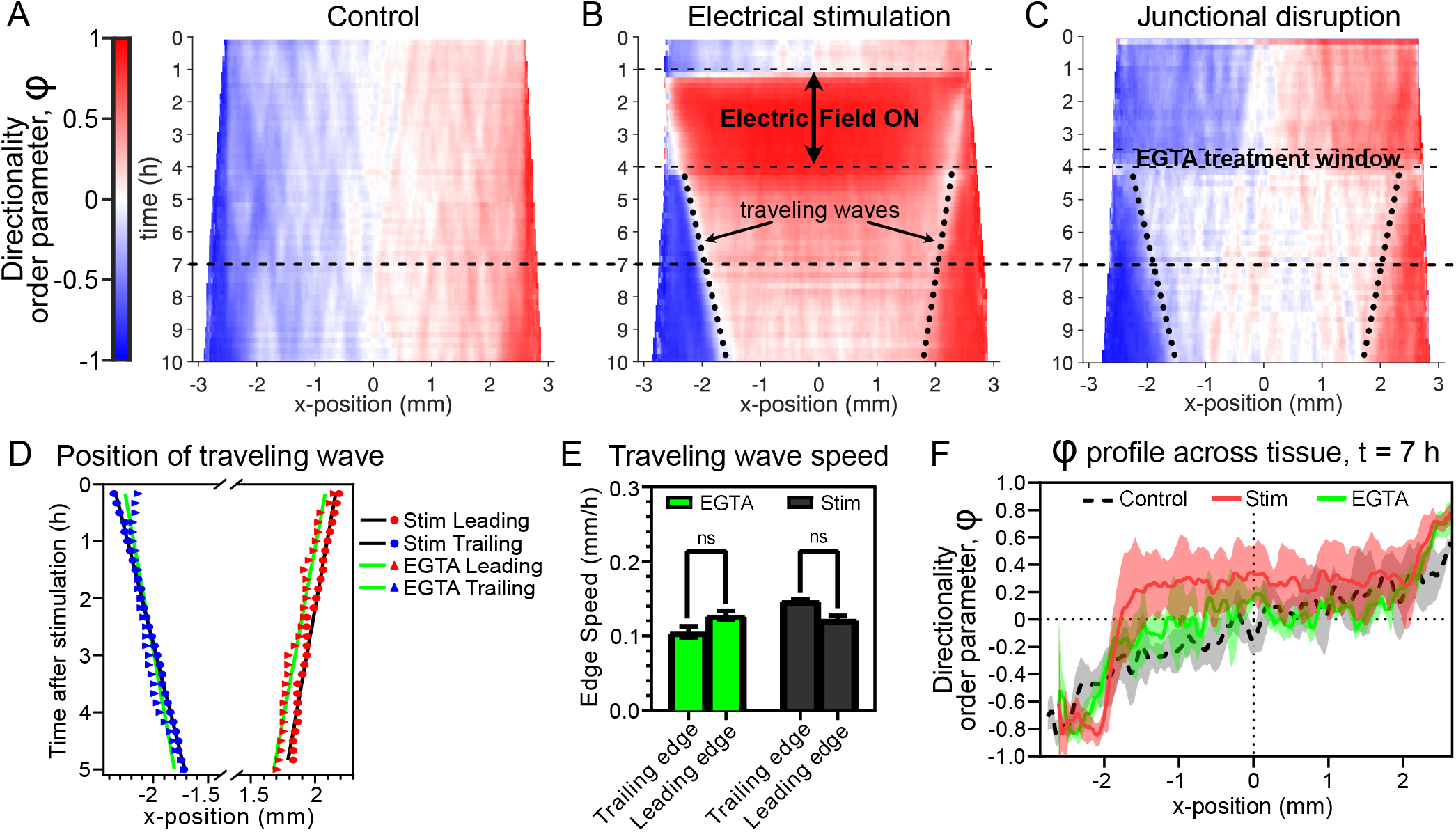
Kymographs reveal apparent mechanical reset post-electrical stimulation. *(A-C)* Averaged directionality order parameter *φ* kymographs (see Methods) demonstrate the evolution of *x*-axis motion patterns across a whole tissue. *(A)* Control tissues, *(B)* electrically stimulated tissues, and *(C)* tissues with EGTA-disrupted cell-cell junctions (see Methods), where the end of treatment was timed to coincide with the end of stimulation in *B* for optimal comparison. Dashes diagonal lines in *B-C* demarcate the ‘triangles’—inward traveling waves of outward migration induced by mechanical reset. *(D)* Plot of the positions of these traveling waves in *B,C*; lines are linear best fits, see panel legend and Methods. *(E)* Traveling wave speeds from slopes of linear fits in *D*: EGTA junction disruption (green); and electrically stimulated tissues (gray). Error bars indicate standard deviation. *(F)* Profile at t = 7 h of directionality order parameter *φ* across all three kymographs, as marked by the dashed black line across panels *A-C*. Shading indicates standard deviation across tissues. For all panels, *N_control_* = 6, *N_stimulated_* = 9, *N_EGTA_* = 3.

We hypothesized that these large-scale inward traveling waves indicated that removal of the global electrical cue significantly altered the mechanical state of the tissue. It has previously been demonstrated that *steady-state* electrotaxis can alter tissue mechanics with increased traction force magnitude in a tissue center and reoriented traction force alignment perpendicular to the electric field vector (54), hence, sudden removal of the electrotactic cue should also alter mechanics. However, a better analog to the large, inward waves we observed comes from a prior study without electrical stimulation that demonstrated that transient disruption of cell-cell adhesion within an epithelium appeared to reset the internal mechanical tension and cell traction state of the tissue, while also inducing similar large-scale inward waves (63). For comparison, we replicated this perturbation using a 30 minute pulse of EGTA (Fig. 4C; see Methods) to briefly chelate calcium and transiently disassemble E-cadherin binding between cells in an otherwise control tissue, and observed remarkably similar inward traveling waves of cell mobilization upon restoration of calcium. We confirmed that the inward traveling waves induced by both electrical stimulation and calcium chelation featured linear propagation at rates of ~ 100 *μ*m/h (Fig. 4D-E), in agreement with those previously described (63) and consistent with a large-scale mechanical ‘reset’ caused by removal of the global bioelectric command.

While halting electrotaxis and transiently disrupting cell-cell adhesion produced similar effects on tissue boundaries, a key difference in the response lies in the bulk tissue behavior, where post-stimulation tissues still exhibited directional memory in the bulk (Figs. 3, center of 4B in 4-10 h range), while chemical disruption of cell-cell junctions largely shut down cell motility in the bulk (center of Fig. 4C in 4-10 h range). We quantified this by analyzing line sections taken across each kymograph at t = 7 h (Fig. 4F), giving us a snapshot of the profile across the tissues 3 h after the respective perturbations. Control tissues exhibited a smooth and largely symmetric, graded directionality profile from the center of the tissue outward (Fig. 4F, black dashed curve). Tissues with transient chemical disruption of cell-cell junctions exhibited highly directed edge zones, but no net directionality in the bulk (Fig. 4F, green curve). However, post-stimulation tissues exhibited not only similar directed edge zones to the junctional disruption case, but a central zone with net positive directionality, implying persistent rightward motion long after stimulation had ceased (Fig. 4F, red curve, matching ‘memory’ shown in Fig. 3). Hence, ending electrical stimulation appears to reset the tissue’s mechanical state at the boundaries similarly to chemically disrupted cell-cell adhesion. However, despite similarity in migratory behavior at tissue edges, it is unlikely that electrotaxis disrupts cell-cell junctions as E-cadherin has been shown to be necessary for MDCK and certain other epithelial electrotaxis (23, 24). Moreover, the migratory memory and preservation of front-rear polarity in the tissue bulk was only observed post-electrotaxis, and not post-junctional disruption.

### Programmed migration with electrotaxis disrupts collective strain wave propagation

As commanding a tissue to migrate ‘rightward’, and abruptly negating that command, both appear to alter the tissue mechanical state in a location dependent fashion akin to junctional disruption, we next sought a biophysical mechanism coupling collective migration behaviors across a tissue. Critically, the waves from Fig. 4 represent coordinated inward-traveling mechanical waves of outward-directed migration, which ultimately allow the motile tissue edges to mechanically influence migration within the tissue bulk. For instance, if a leading edge cell in a cohesive tissue migrates outward, then its immediate rearward-adhered neighbor can follow, allowing that cell’s own rearward neighbor to follow suit, and so on, thus forming a wave of force-coupling and cell mobilization. Such waves have been shown to create zones of stretching and compressing, that are transmitted across the tissue via ‘strain waves’, and tend to originate at tissue boundaries (57, 63). To capture these waves and analyze the overall rate of cellular deformations, we measured the *rate* of strain—stretching (positive) or compressing (negative)—from the PIV vector fields, as 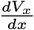 for strain rate in the *x*-direction 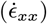 and 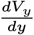 for strain rate in the *y*-direction 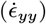.

Since strain waves are difficult to visualize in such large tissues, we represented our control tissue bulk data as strain rate kymographs, a horizontal strip for 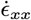 (Fig. 5A) and a vertical strip for 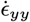 (Fig. 5B), from the strain rate vector fields, similar in form and presentation to the strain waves depicted in prior studies (63). Waves of 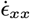 traveled primarily in the *x*-direction, appearing as diagonal regions of stretch (tensile strain rate; purple in Fig. 5A) or compression (compressive strain rate; green in Fig. 5A) in the *x-t* kymograph, with the slopes confirming that these waves propagated at similar rates as the triangular waves previously discussed in Fig. 4. Similarly, 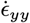 waves traveled primarily in the *y*-direction, seen in the *y-t* kymograph (Fig. 5B, also depicted with sample slope speed illustrations). Movie 5, with control and stimulated tissues side-by-side, visually portrays general changes in strain rate dynamics, especially the boundary and bulk effects post-stimulation.

**Fig. 5.**
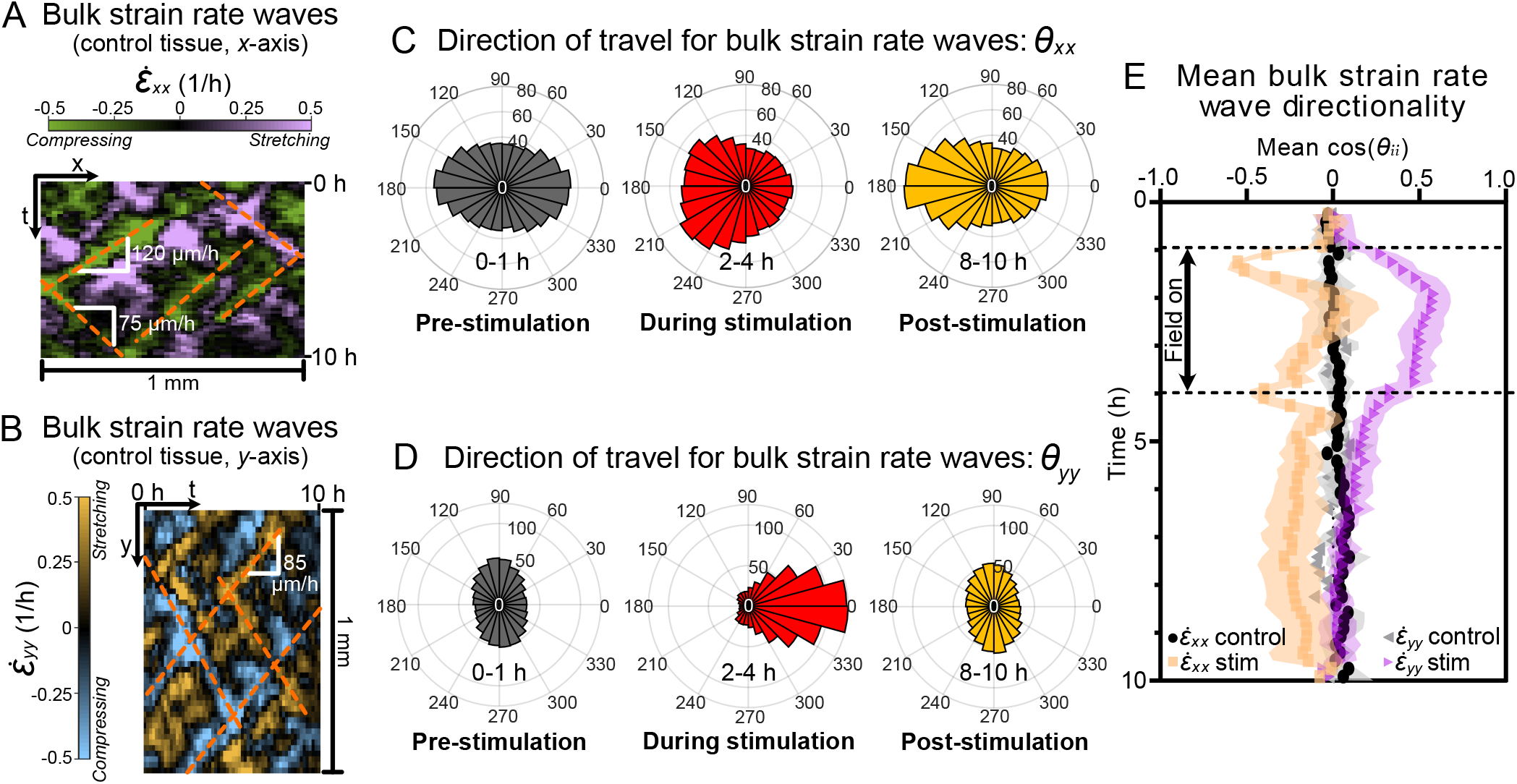
Strain waves reorient based on electrical stimulation. *(A)* Representative *x-t* kymograph (1 mm over 10 h) of bulk strain rate 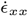 in unstimulated tissues showing horizontal strain wave propagation. Positive strain rate (purple) indicates stretch; negative strain (green) indicates compression. Wave propagation speeds (slope of dashed orange lines) varied from 75 *μ*m/h to 120 *μ*m/h. *(B)* Representative *y-t* kymograph of bulk strain rate 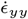; note vertical orientation relative to *5A* to emphasize vertical strain wave propagation. *(C)* Polar histograms of strain rate wave direction, *θ_xx_*, in stimulated tissues plotted for three experimental regimes: unstimulated (gray, 0-1 h); stimulated (red, 2-4 h); post-stimulation (yellow, 8-10 h). Note sustained leftward shift of *θ_xx_* post-stimulation. *(D)* Polar histograms of strain rate wave direction, *θ_yy_*, analogous to *5C*. Note transient rightward orientation during stimulation (red), followed by recovery to baseline post-stimulation (yellow) strongly matching the unstimulated case (gray). *(E)* Quantification of average strain wave directionality (mean *cos*(*θ_ii_*); −1 is leftward, +1 is rightward) for each condition. Control tissue cases of 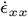 (black) and 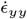 (gray) show no net directionality. Orange squares show mean *cos*(*θ_xx_*) time-course in stimulated tissues. Note perturbation during stimulation and persistent lack of recovery, biased leftward. Purple triangles show mean *cos*(*θ_yy_*) in stimulated tissues. Note rightward perturbation during stimulation and clear recovery post-stimulation. Shading indicates standard deviation across tissues. Panels *C-E* data is averaged over N=9 stimulated tissues.

Kymographs of strain rate only visually capture wave propagation that persists for several hours. To quantify the propagation direction of these waves in the shorter timescales of our experiments, we performed additional PIV analysis on image sequences of strain waves themselves (see Methods). This produced vector fields of the primary propagation direction of strain waves at each location within the tissue for every timepoint, which we denote as *θ_xx_* (for 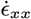 waves) and *θ_yy_* (for 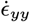 waves). We then plotted polar histograms of *θ__xx__* and *θ_yy_* within the tissue bulk to visualize the distribution of strain rate propagation direction (Figs. 5C-D). In the unstimulated control hour (0-1 h; gray-shaded left histograms of Figs. 5C-D), as in control tissues (above, Figs. 5A-B), 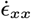 waves travel primarily horizontally with equal leftward- and rightward-moving waves (Fig. 5C, left), and 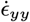 waves travel primarily vertically with balanced upward- and downward-moving waves (Fig. 5D, left). This matches expectations whereby information in a symmetric, unperturbed tissue should propagate equally from all free edges throughout the tissue.

Surprisingly however, these dynamics dramatically changed during and after electrical stimulation, where the 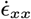 strain waves primarily propagated leftward, with this effect being even more pronounced post-stimulation than during electrotaxis itself (Fig. 5C, center and right). This means that globally ‘rightward’ electrotaxis induces a mechanical tissue state, such that horizontally-traveling waves within the tissues are dominated by those that would normally stem from the rightward (leading) edge. By contrast, 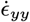 waves were strongly biased in the direction of stimulation during electrotaxis (90 degrees of their baseline orientation), essentially shifting with the cell migration itself, but this effect relaxed post-stimulation to match control orientation (Fig. 5D, center and right). These data suggest that, while active electrotaxis appears to reprogram much of the endogenous mechanical strain state within a tissue, the lasting impact lies only along the axis of induced migration.

To further illuminate this, we analyzed the dynamics of the strain wave disruption process using the average value of *cos*(*θ_xx_*) and of *cos*(*θ_yy_*) within the tissue bulk as a metric for the directionality of 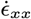 and 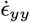 waves, respectively (Fig. 5E). A directionality of 1 would represent a tissue for which waves are only traveling parallel to the field, while a directionality of −1 would represent waves only traveling antiparallel. 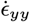 directionality peaked to the right about 1 h after stimulation was initiated and decayed quickly after stimulation was turned off (Fig. 5E, purple triangles), a time-course which is strikingly reminiscent to that of speed (Fig. 3B). 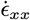 waves were more nuanced, with large negative jumps in directionality in the first timepoint of stimulation and the first timepoint post-stimulation, each followed by dynamic changes which are perhaps tied to viscoelastic effects. Post-stimulation, however, the 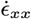 directionality eventually reaches a distinct new steady state—a long-term bias toward the trailing edge with no apparent decay (Fig. 5E, orange squares). The initial immediate changes to wave propagation bear resemblance to waves of signaling that span a tissue in the matter of minutes as a response to tissue damage, which also promptly move leftward from a wound at the right edge (65–67), in contrast to the much slower mechanical waves that produced the boundary ‘triangles’ in the kymographs in Figs. 4B-C and of migration upon release of a barrier (63). In these data, we see that both initiation *and removal* of stimulation strongly reprogrammed the mechanical state of the tissue. This long-lasting change of mechanical communication should produce changes in the collective structure of cell migration, which we analyze next.

### Driving collective cell migration overrides and weakens cell– cell crowd interaction dynamics even post-stimulation

A key hallmark of any collective motion process is that information couples well beyond simple nearest-neighbor interactions. In bird flocks, neighbor correlation dynamics may propagate up to 7 neighbors (27), while the MDCK epithelium is known to exhibit correlated domains of 5-10+ cells (68, 69), mediated by cell-cell adhesion (70–72). However, programming a particular pattern of migration into a group that already possesses internal collective coupling necessitates a conflict (7), mediated in electrotaxing epithelia by E-cadherin cell-cell adhesion (4, 7, 23). Given this framework, we hypothesized that electrically programming collective cell migration might reprogram the endogenous correlation length to force cells to entrain to the migration command, and that the relaxation behaviors we observed were driven by a re-establishment of the native correlation dynamics.

To test this hypothesis, we calculated the velocity-velocity spatial correlation length both parallel (*V_x_*) and perpendicular (*V_y_*) to the electric field stimulation axis at each time point using PIV data. The correlation length reflects the approximate size of correlated domains within a larger population. When calculating velocity-velocity correlations in a highly directed system (e.g. flocks of birds, road traffic, or ensembles of electrotaxing cells), it is critical to first subtract the global mean velocity at any given time and therefore perform correlation analysis on the velocity residuals, or fluctuations about the mean, which much better capture the internal cell-cell interactions by removing the global bias (39, 73). This effect is demonstrated in Figs. 6A-B, where we compare the raw velocity field to the residuals, respectively, with insets that emphasize the importance of correlating the residuals in our directed migratory system. The averaged correlation over time and its statistical comparisons are shown in Figs. 6C-D. Representative correlation curves are shown in Fig. S4, and our approach is described in Methods.

**Fig. 6.**
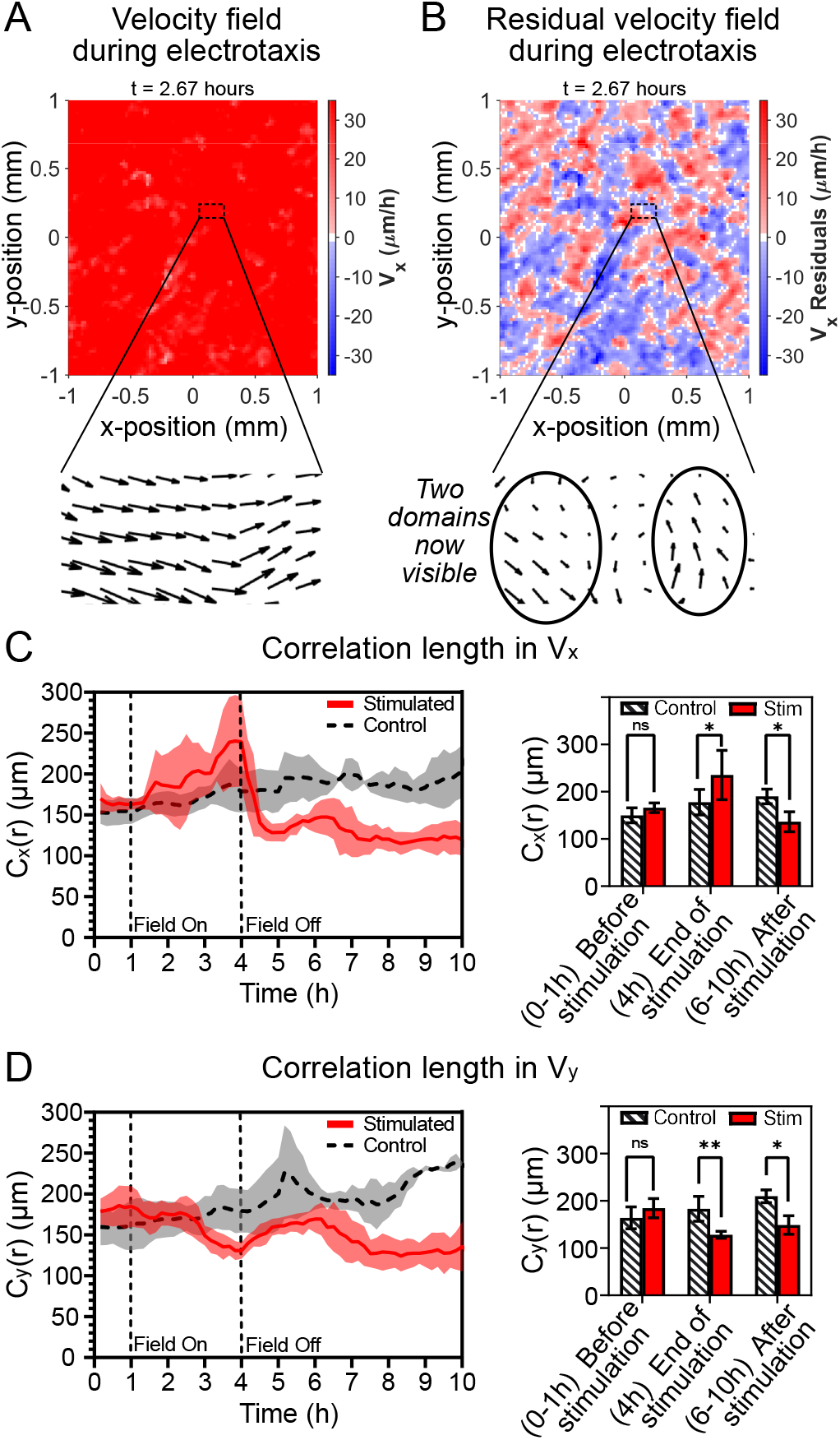
Electrical stimulation has a lasting effect on spatial correlations. *(A)* Velocity heatmap during electrotaxis showing the global stimulus response of the central 1 × 1 mm region of a representative tissue. Inset shows velocity vector field in a 200 × 100 *μ*m zone. *(B)* Corresponding heatmap of velocity residuals during electrotaxis emphasizing many small, correlated domains. Corresponding inset to *A* showing underlying domains. *(C),left* Temporal dynamics of correlation length for *V_x_*: *C_x_* (*r*) showing increased correlation during stimulation and decreased correlation post stimulation. *(C),right* Statistical comparisons of *C_x_* (*r*) relative to control (*, *p* = 0.04) emphasize ~40% relative increase during stimulation and ~40% decrease post-stimulation. *(D),left* Temporal dynamics of correlation length for *V_y_*: *C_y_* (*r*). *(D),right* Statistical comparisons of *C_x_* (*r*) relative to control (**, *p* = 0.004; *, *p* = 0.04). Note sustained drop in correlation relative to control by 10 hrs. For all correlations, shading and error bars indicate standard deviation; *N_stimulated_* = 9, *N_control_* = 6.

We first analyzed correlation length parallel to the field axis, *C_x_* (*r*), as shown in Fig. 6C. For reference, we initially quantified the *x*-axis velocity correlation length of control tissues over the experimental time period (Fig. 6C, dashed black curve), confirming the expected gradual increase over time (61). By contrast, during stimulation, we noted a significant increase (nearly 40% by the end of stimulation; Fig. 6C red curve, and center of bar plot) compared to unstimulated control tissues of *C_x_* (*r*), followed by a similarly strong reduction relative to control tissues after the stimulus was removed (Fig. 6C, right side of bar plot). However, unlike our other metrics such as the directionality order parameter or mean velocity (e.g. Figs. 2,3), *C_x_* (*r*) showed no signs of equilibrating to the control tissue behavior, even 6 hours post-stimulation—twice the stimulation period (Fig. 6C, plateau trend from 7 h on, and right side of bar plot). As mechanical strain state exhibited different behaviors parallel and perpendicular to the field axis, we also analyzed how correlation lengths changed perpendicular to the field axis, *C_y_* (*r*) (Fig. 6D). Here, we saw a significant decrease in correlation length (approx. 25%, Fig. 6D red curve, and center of bar plot) relative to the control tissue by the end of the stimulation period, followed by similar behavior to what we observed in *C_x_* (*r*) post-stimulation, where *C_y_* (*r*) again remained significantly depressed relative to correlations in an unstimulated control tissue by the tenth hour (Fig. 6D, plateau trend from 7 h on, and right side of bar plot).

These data emphasize several key details. First, there appears to be a trade-off between *C_x_* (*r*) and *C_y_* (*r*) during stimulation, where stronger correlations of cell migration fluctuations along the field axis necessitate weaker correlations in the orthogonal axis. More surprising, perhaps, was the clear reduction in correlation length that occurred in all axes post-stimulation relative to a control tissue (right sides of bar plots in Figs. 6C-D). This reduction in correlated domain size again implies that the act of removing the field stimulus appears to trigger a long-term, large-scale loss, or a ‘reset’ of the tissue state relative to an unperturbed tissue—consistent with the resetting of biophysical behaviors shown in Fig. 4. Further, this emphasizes that our earlier hypothesis was incorrect—the post-stimulation speed and directionality relaxation dynamics do not, in fact, reflect a restoration of baseline correlations.

## Discussion

This study furthers our understanding of bioelectric manipulation of collective cell migration and general control of group dynamics in two key regards. First, we comprehensively assessed the complete step response of a tissue undergoing electrotactic migration, from onset of directed migration to long-term relaxation. Next, we linked these temporal dynamics to specific, supracellular patterns of electrotaxis at the tissue scale. Together, these results revealed both that short-term driven migration can produce longer-term changes to collective cell behaviors (e.g. Figs. 3,4), and that these effects play out differently throughout the tissue (e.g. bulk, leading, trailing, or perpendicular edges) as in Figs. 1,2,5. More broadly, our data imply that globally driven directed cell migration (mediated by electrotaxis here) appeared to ‘reset’ key mechanical aspects of the stimulated tissue, both during and long after stimulation, affecting not only collective tissue growth dynamics (Fig. 4), but also cell-cell interaction range (Fig. 6), apparently mediated by alterations to how mechanical strain coupled across cells (Fig. 5).

How migratory programming affects a tissue in time is at least as important as the spatial, or supracellular response, as large variations in response timescales contributed to the overall emergent collective behaviors during and after stimulation. Many tissues, such as epithelia, are tightly cohesive due to cell-cell interactions, which gives rise to both elastic and viscous tissue mechanics at various timescales (74–76). In our stimulated tissues, rapid responses (<1 hr) include the sharply recoiling edge zones (Fig. 2), the short post-stimulation speed equilibration (Fig. 3), and the quickly propagating and shifting parallel strain waves during and after stimulation (Fig. 5E). These fast biomechanical responses contrast with the multi-hour equilibration periods after stimulation, such as the sustained rightward speed drop in recovering edges (Fig. 2C), the very slow reversion to undirected migration in the tissue bulk (Figs. 3A,C,E), and the return to coordination through strain waves perpendicular to the field (Figs. 5D-E). Perhaps most surprising was that coordination through strain waves parallel to the field and cell-cell correlations in the bulk of the tissue (Figs. 5D,E; Figs. 6C,D) displayed no sign of recovery during the entire post-stimulation period—a regime at least twice as long as the initial stimulation. These are reminiscent of the different timescales in response to mechanical stretch (74–76), and while electrotaxis does not necessarily stretch the footprint of the tissue as a whole (54), it does induce displacement and internal deformation by directing tissue flow differently from that observed in an unperturbed tissue. As we have shown that this flow plays out in a tissue with supracellular behavioral zones, it should necessarily cause longer term plasticity of tissue properties as each zone behaves differently from each other. Future studies integrating these results with whole-tissue traction force and cell-cell force analysis would help to clarify this.

Such plasticity and susceptibility to control, along with the apparent mechanical reset (Fig. 4), might explain why the tissue maintains such a strong memory of directionality in the tissue bulk (Fig. 3) post-stimulation, as the underlying mechanical state had been entrained to the command direction. If each and every cell can sense the electrical stimulation, the global command could eliminate the need for tissues to coordinate flow with neighbors. While single-cell mechanical analysis would be needed to better explore this, we saw clues in our correlation length analysis (Fig. 6). In the direction of stimulation, fluctuations around the mean velocity undergo an increase in length scale during electrotaxis—implying longer-range cell-cell coordination—which may be necessary to maintain tissue integrity at 2-5× increased migration speeds induced by electrotaxis (Fig. 3B). Further, the coordination realized during electrotaxis must be qualitatively different than that of standard tissues as it is asymmetric (Fig. 6C vs. Fig. 6D) and is quickly lost after stimulation is removed; this is consistent with previous work showing that the tissue tension profile aligns perpendicular to the stimulation direction (54).

More generally, that even a simple, universal ‘command’ produced such nuanced behavior highlights not only the importance of taking into account supracellular, or supracollective regional behaviors when controlling groups but, perhaps most importantly, that these behaviors may also differ from each other and from expectations post-stimulation. Specifically, our data clearly demonstrated that turning off a stimulus does not simply allow a collective system to return to nominal behavior, meaning that stopping stimulation can be just as transformative to tissue or group dynamics as starting it. These concepts are important to consider in any collective system, whether trying to control theoretical active fluids (77), to steer cells (18, 35), or to herd sheep (78). In the longer term, understanding the relevant timescales and coupling behaviors in driven collective motion will likely help to improve stimulation and control efficacy. Understanding the long-term effectiveness of short-term migration stimulation in a given system can help reduce the need for long term or more complex stimulation— especially valuable in contexts such as *in vivo* applications or therapeutics, including bioelectric bandages to accelerate wound healing (22). Overall, these approaches and concepts should also provide a foundation for comparisons of different classes of global commands in tissues (e.g. chemotaxis or optogenetics), as well as other collective systems. Finally, we suspect that more optimal control in the future will involve more tailored ‘commands’ specific to each functional region within a group.

## Supporting information

SI Appendix - Wolf et al 2021

Movie 1 - full tissue phase time-lapses

Movie 2 - phase 0.5mm strips time-lapses

Movie 3 - velocity heatmaps

Movie 4 - top zone tracks

Movie 5 - strain wave heatmaps

## Data Availability Statement

High resolution, raw data are available on a Zenodo repository (10.5281/zenodo.5120700). Relevant codes from our analyses are available on Github at github.com/CohenLabPrinceton/ElectrotaxisSupracellularMemory

## Materials and Methods

### Cell culture

MDCK-II wild type canine kidney epithelial cells were a gift from the Nelson Laboratory at Stanford University and were cultured in customized media consisting of low-glucose (1 g/L) DMEM with phenol red (Gibco, USA; powder stock), 1 g/L sodium bicarbonate (lower than standard DMEM), 1% streptomycin/penicillin (Gibco, USA), and 10% fetal bovine serum (Atlanta Biological, USA). Cells were maintained at 37°C and 5% CO_2_ in humidified air.

### Micropatterning of epithelial arrays

Standardized arrays of epithelia were produced using silicone tissue stencils built by razor-writer as described previously (18, 33). A brief summary follows. 250 *μ*m thick sheets of silicone elastomer (Bisco HT6240) were cut into stencil patterns (5 × 5 mm square arrays) using a razor writer (Silhouette Cameo) and transferred into culture vessels. Suspensions of MDCK cells were then seeded into the stencil patterns with the volume and density tuned to produce uniformly dense tissues. Three ranges of cell densities were analyzed here as follows. Medium density tissues, averaging 2740 ± 320 (SD) cells/mm^2^, were our baseline samples and intentionally covered a large range. Tissues classified as high density tissues averaged 4450 ± 90 (SD) cells/mm^2^ and exhibited limited movement at the start of the experiment, as was expected (58). Low density tissues, averaging 2230 ± 90 (SD) cells/mm^2^, were at the lowest density while still maintaining confluence in the 5 × 5 mm space. In all cases, cells were cultured for 1 h to allow for adhesion and then the dish was flooded with media and maintained for 16 h in an incubator prior to being inserted in our electro-bioreactor as described below.

### Electro-bioreactor Design

The electro-bioreactor design used here was modified from our SCHEEPDOG platform (18) for uniaxial stimulation over a large culture area as described previously (7). Briefly, a custom laser-cut acrylic housing, combined with silicone adhesive layering for a tight seal, was placed around a microarray of tissues in a 10 cm tissue-culture dish to allow media perfusion from north to south and electrical stimulation from west to east. The electro-bioreactor accommodated a 35×17 mm cell channel, flanked on both sides by salt bridges of 4% w/v agarose in phosphate-buffered saline (PBS), through which the electric current passed from Ag/AgCl electrodes, each sitting in a reservoir of 2 mL PBS. Titanium probes were inserted into the agarose salt bridges to monitor the voltage across the channel throughout the assay; by connecting these to a USB oscilloscope (Analog Discovery 2, Digilent Inc.), we were able to finely-tune the electric current sourced by a Keithley 2450 SourceMeter (Tektronix) with proportional feedback control using a custom MATLAB script to ensure 3 V/cm during stimulation and no current otherwise. For a complete summary of the approach and guidelines, see Zajdel et al (18) and our repositories (github.com/cohenlabprinceton).

In this study, every electrotaxis experiment began by removing the stencil surrounding the tissues 2 h before data collection. After a 1 h unstimulated control period, electrical stimulation was induced at 3 V/cm across the channel for 3 h before the field was removed while imaging continued for the remainder of the experiment. For the assay with chemical junctional disruption, epithelia were otherwise treated as control tissues, but the perfusion media fed into the electro-bioreactor was doped with 4 mM of EGTA (EMD Millipore Corp., USA) for 30 minutes, from the 3.5 h to 4 h mark, timed to end at the same time as the end of stimulation in our electrotaxed tissue experiments.

### Time-lapse imaging and data collection

We captured timelapse microscopy images every 10 min with an automated inverted microscope (Zeiss Axio Observer Z1) equipped with an XY motorized stage, controlled using Slidebook (3i Intelligent Imaging Innovations). The microscope was equipped with a 5x/0.16 phase contrast objective, a Photometrics Prime (Photometrics, Inc.) sCMOS camera, and a custom-built cage incubator to maintain 37°C. Inside, fresh media with continuously bubbled 5% CO2 was perfused with a peristaltic pump (Instech Laboratories) through the electro-bioreactor at a rate of 2.5 mL/h.

### Image post-processing, particle image velocimetry (PIV), and nuclear cell tracking

FIJI (https://imagej.net/software/fiji) was used to process all tiled time-lapse images through template-matching (79), stitching (80), and masking. Velocity vector fields were calculated from the processed time-lapse images using particle image velocimetry based on PIVlab (81, 82), with a two-pass iteration of 64×64 pixel and 32×32 pixel interrogation windows, both with a 50% step size, providing a 16-pixel final step size between vectors. We filtered out vectors that were outside five standard deviations and replaced them with interpolation. All data was then analyzed with MATLAB (Mathworks, 2019a), providing visualizations of the cell movements across the entire tissue.

Cell nuclear and density data were calculated by segmenting images from our in-house Fluorescence Reconstruction Microscopy tool (83), a convolution neural network trained to produce images of nuclei from our phase contrast images. We also used this output to calculate trajectories of cell nuclei within a 150 *μ*m thick zone from the edges, taking only the center 2 mm near each edge. We linked cell tracks with Linear Motion Tracking, after detecting spots filtered by the Laplacian of the Gaussian, using FIJI’s TrackMate plugin (84).

### Average kymographs

We first created representative kymographs by averaging over the vertical direction across the entire width of each tissue, excluding the outer 1.5 mm of the top and bottom. We then aligned kymographs based on the initial centroid of each tissue and averaged across all tissues for each condition. We did not plot points at the outer boundary that had not yet been reached by more than a single replicate.

### Strain wave propagation analysis

PIV analysis requires grayscale images, so we mapped 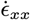 and 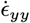 vector fields from all timepoints to 8-bit grayscale image sequences. As such, we mapped the maximum and minimum 0.1% of strain rate values from each vector field to 255 and 0, respectively, interpolating the remaining values to the appropriate integer between 0 and 255. We then performed PIV analysis on the resulting image sequences, using 32×32 followed by 16×16 pixel windows with 50% overlap. Note that the resulting fields of strain rate propagation will then have resolution 8x less than strain rate. Each pixel of strain propagation then represents a 256×256 window of the original phase contrast images.

### Characteristic time scale analysis

Characteristic time scales were obtained by fitting exponential decay curves to mean |*V_x_*| and directionality order parameter *φ* data using a custom MATLAB script. A nonlinear model was fit to the data after the end of stimulation, from the fourth through tenth hour, using the model functions 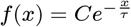 for mean directionality order parameter and 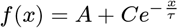 for mean |*V_x_*|. Initial values for the model were taken directly from the data. R^2^ was greater than 0.91 for all fits.

### Correlation length analysis

Spatial correlation lengths were calculated using a custom MATLAB script. Briefly, 2-D spatial autocorrelation was performed on PIV vector field data utilizing the xcorr2() function, and a radial scan of the autocorrelation matrix was used to obtain a correlation curve. The correlation length was obtained by interpolating the correlation curve and calculating the distance at which the correlation drops below 10% of its maximal value.

### Traveling wave analysis

Inward traveling waves of cell mobilization visualized in mean directionality order parameter kymographs of both electrically stimulated and EGTA-treated (chemical junctional disruption) tissues alike were analyzed using a custom MATLAB script. Kymographs were trimmed and binarized before masked using a watershed. Regions of the image were then filled and closed boundaries were located using built-in morphological operations. The traveling waves were then found by finding the first non-zero pixel in each respective half of the image. For speeds, a linear fit was made to the discovered waves, and slopes were reported.

### Statistical analysis

Statistical tests were conducted using GraphPad Prism 9.1.2 (GraphPad Software) with an unpaired two-sided Mann-Whitney non-parametric U test. When comparing a variable between stimulated and control tissues, ensemble values (and not, for example, individual PIV vector data) for each tissue were calculated and then each classification group was compared one against the other.

## ACKNOWLEDGMENTS

Support for this work was provided in part by National Institute of Health Award R35 GM133574-03 and National Science Foundation CAREER Award 2046977. We thank the members of D.J.C’s lab for feedback and support, and Gawoon Shim for help with bioreactor device set-up.

