## Supplementary material for "Short-term stimulation of collective cell migration in tissues reprograms long-term supracellular dynamics": SI Appendix - Wolf et al 2021

**This PDF file includes:**

Figures S1 to S4
Legends for Movies 1 to 5

**Other supplementary materials for this manuscript include the following:**

Movies 1 to 5

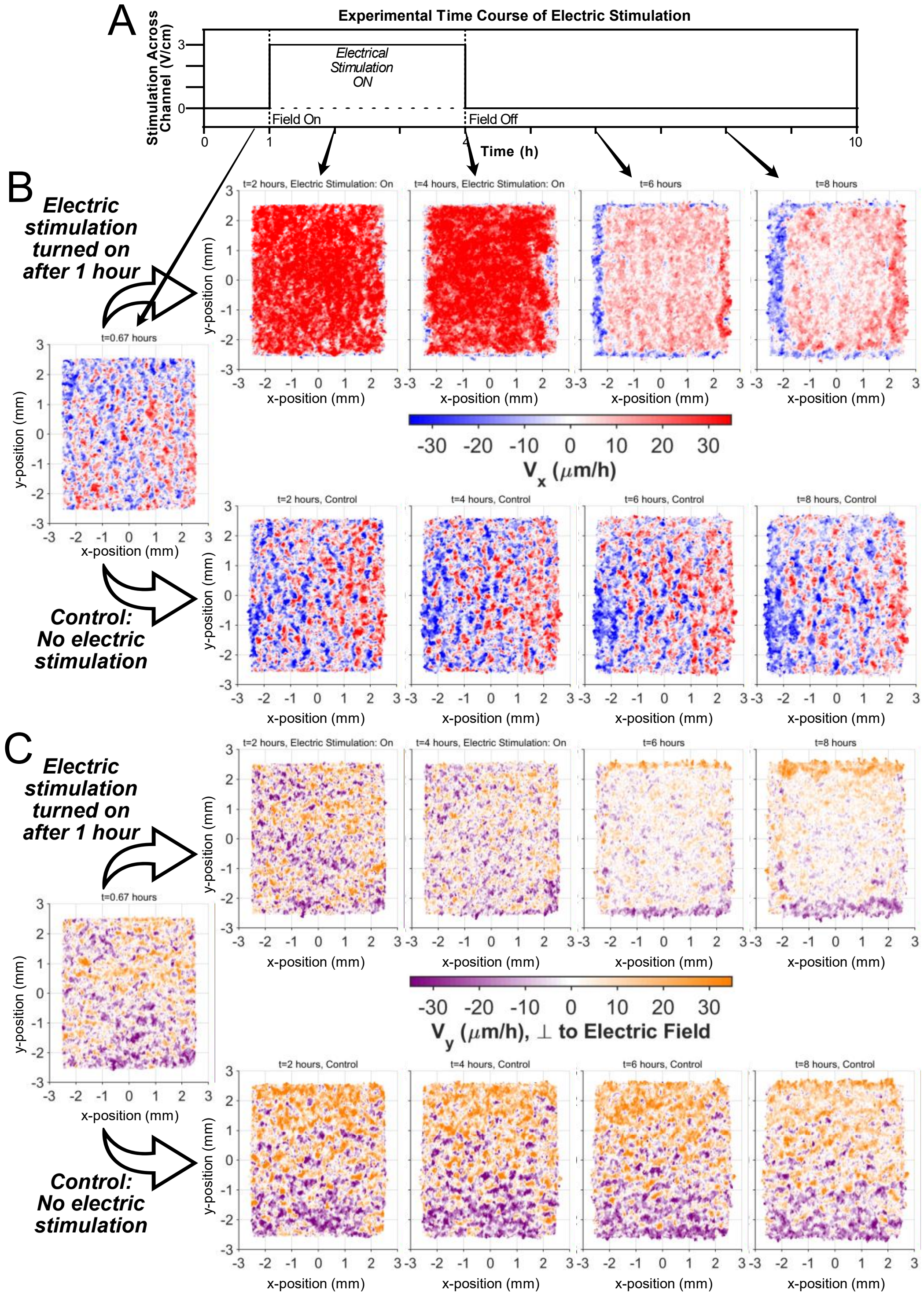


**Fig. S1.** $V_x$ and $V_y$ heatmaps for stimulated and control tissues. **(A)** Time course of our experimental setup, to see the complete step-response to bioelectric stimulation: 1 h of pre-stimulated control time; 3 h of stimulation ON; and 6 h of relaxation with stimulation OFF. **(B)** $V_x$ heatmaps for stimulated and control tissues throughout the experiment. Stimulated tissue heatmaps showcase global directed motion during stimulation (strong red color), and retrograde motion in the edges after stimulation ends, with a pink center displaying 'memory' in the tissue bulk. **(C)** $V_y$ heatmaps for stimulated and control tissues throughout the experiment. Stimulated tissue heatmaps have no global directed motion as seen in control tissues, however after stimulation ends, most of the motion is focused in the edge regions.

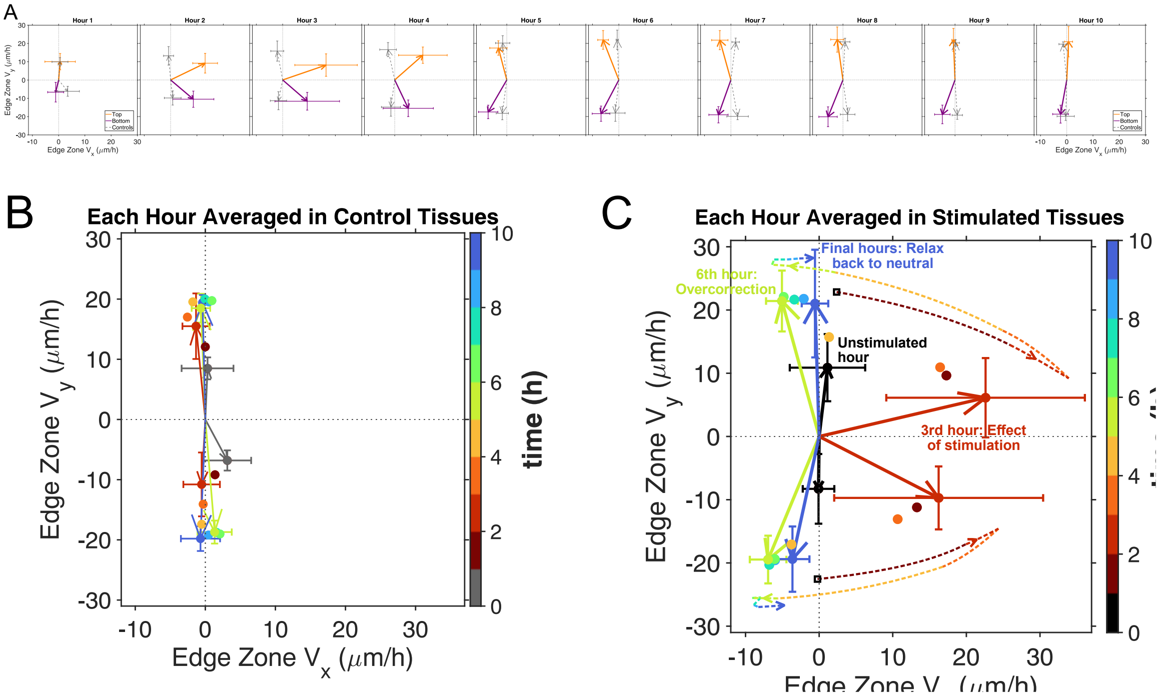


**Fig. S2.** Mean velocity vectors in the 150 µm-tall zones near the top and bottom edges show unique dynamics during and after stimulation. Stimulated tissue vectors exhibit similar behavior as the control before stimulation begins (first hour), obvious rightward motion during stimulation, then recoil opposite the direction of stimulation after it is turned off. **(A)** has 10 panels, each representing the average across the entire respective hour during the experimental time course. Electric stimulation starts after the first panel, and ends after the fourth. In panels 2-4, the top (orange) and bottom (purple) average vector is strongly biased with the direction of stimulation. Immediately after, however, in panel 5, the vectors sharply flip back, overcorrecting toward the left. This overcorrection slowly relaxes over time through the 6 h post-stimulation period. Control vectors are provided in gray. **(B)** These same vectors are overlaid for the control tissue. Four vectors are shown, with the other six represented only by a filled-in circle. **(C)** Similar to panel B, but stimulated tissue data. Dashed lines and annotations were added to help visualize the development of the average vector over time, during the three phases of our experiment as described above. For all panels, all data was averaged over N=9 for stimulated tissues, N=6 for control tissues, and within a 150 µm-tall zone that excluded 1.5 mm on both the left and right for edge effects.

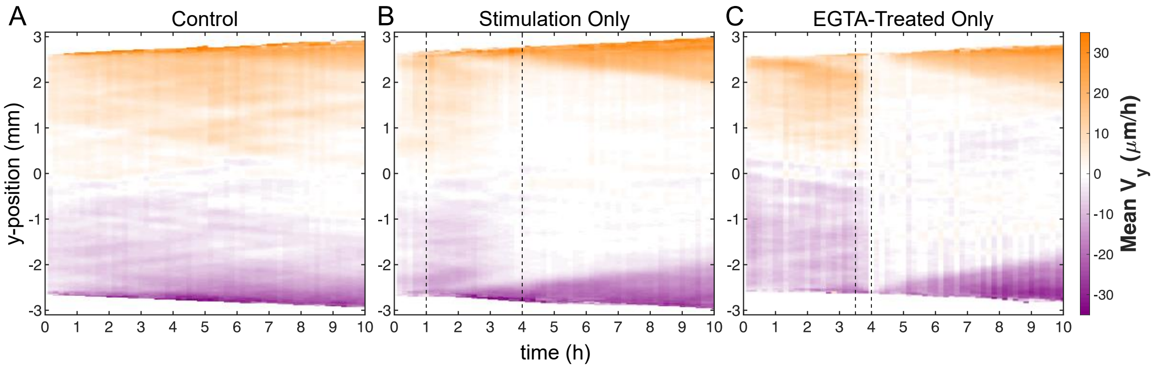


**Fig. S3.** Average kymographs across several tissues of mean $V_y$ in **(A)** control (N=6), **(B)** stimulated (N=9), and **(C)** EGTA-treated (chemical junctional disruption, N=3) tissues. After their respective perturbations have concluded at t = 4 h, in both stimulated and EGTA-treated tissues, inward traveling waves of cell mobilization are clearly visible, just as demarcated in Figs. 4B-C for directionality order parameter kymographs. To calculate mean $V_y$ across the entire height of the tissue, the vertical velocity vector components were averaged at each y location through the x-center of the tissue, ignoring 1.5 mm on the left and right for edge effects.

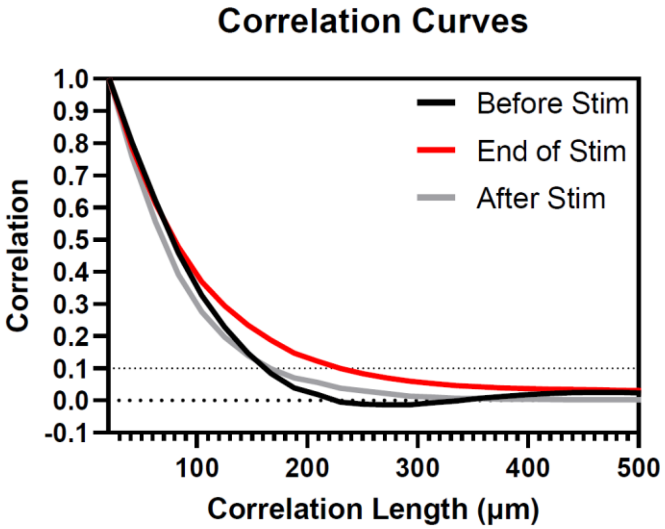


**Fig. S4.** Correlation curves at different periods in the experiment averaged over all stimulated tissues. The correlation curve at the end of stimulation (red) drops much slower than those before (black) and after (gray) stimulation, denoting a higher correlation length within the bulk of the tissue. (See Methods.)

**Movie 1 (separate file). Phase-contrast time-lapse images of 5 x 5 mm MDCK tissues**, side-by-side: a control tissue on the left, and an electrically stimulated tissue on the right undergoing 1 h of control, 3 h of electrical stimulation to the right, and 6 h of unstimulated relaxation.

**Movie 2 (separate file). Phase-contrast time-lapse images of 0.5 mm-tall strips** through the center of the two tissues which appear in Movie 1, providing a sharper video of the dynamics through the entire width of the tissues. Here, the strip of the control tissue appears on top, through its 10 h of unperturbed growth, and the strip of the electrically stimulated tissue appears on bottom, through the three phases of the experiment—1 h of unstimulated control time, 3 h of electrical stimulation 'rightward', and 6 h of relaxation time post-stimulation.

**Movie 3 (separate file). Heatmaps of velocity** parallel to ($V_x$) and orthogonal to ($V_y$) the direction of stimulation through our entire timelapse. The data here is garnered from PIV data (see Methods) for the same two tissues which appear in Movie 1. 10 h of a control tissue appear in the top two panels, and the electrically stimulated tissue appears in the bottom two panels, again through 1 h unstimulated, 3 h electrical stimulation to the right, and 6 h relaxation post-stimulation.

**Movie 4 (separate file). Time-lapse of a sample portion of the top edge of a stimulated tissue, with sample TrackMate tracks** superimposed to highlight the apparent recoil in this area of the tissue. The image itself was generated using our in-house Fluorescence Reconstruction Microscopy tool (see Methods).

**Movie 5 (separate file). Strain wave propagation in both directions for control and stimulated tissues.** In control tissues in the top two panels, strain waves are seen natively moving throughout the tissue; in stimulated tissues however, a particularly stark wave traveled inward from each edge, beginning at the end of stimulation—on the right (leading) tissue edge in x-strains and the top and bottom tissue edges in y-strains. Additionally, in the bulk of the y-strain panel, careful inspection shows strain waves during stimulation moving rightward, even as they propagate vertically, as discussed in the main text with Figs. 5D-E.
